# Mbd3/NuRD is a Key Inhibitory Module During the Induction and Maintenance of Naïve Pluripotency

**DOI:** 10.1101/013961

**Authors:** Asaf Zviran, Yoach Rais, Nofar Mor, Noa Novershtern, Jacob H. Hanna

**Affiliations:** The Department of Molecular Genetics, Weizmann Institute of Science, Rehovot 76100, Israel.

## Abstract

Our group has published a study on induced pluripotent stem cell (iPSC) reprogramming (Rais et al. Nature 2013^1^) that reached the following conclusions: a) Mbd3/NuRD is a repressor of inducing naïve pluripotency from mouse Epiblast stem cells (EpiSCs), primordial germ cells (PGCs), murine somatic cells and human secondary fibroblasts; b) Up to 100% iPSC formation efficiency can be achieved via optimized Mbd3/NuRD depletion, in concert with optimized OKSM delivery and naïve pluripotency conditions (2i supplement applied only after 48 hours, human LIF, hypoxia and Vitamin C containing Knockout serum replacement)^1^. This represented the first proof for deterministic/near-deterministic iPSC reprogramming, and highlighted a previously unappreciated role for Mbd3/NuRD in hampering the re-establishment of pluripotency. Recent reports have seemingly provided contradictory results and attempted to dispute our iPSC efficiency quantifications and/or the role of Mbd3/NuRD in blocking reprogramming^2,3^. Here we provide a detailed response to these reports based on extended discussions and providing new data. The synthesis presented herein disagrees with claims made by Silva, Hendrich, Bertone and colleagues^2,3^, and reconfirms that Mbd3/NuRD is a major pathway that inhibits the maintenance and induction of pluripotency^1^. Further, we foresee that its controlled manipulation is likely to become an integral pathway for inducing and maintaining naïve pluripotency in a variety of species.

## Introduction

Cellular reprogramming has boosted a major revolution in the field of stem cell research^4^. This is a relatively simple process in which the induction of exogenous transcription factors (classically Oct4, Sox2, Klf4 and c-Myc (abbreviated as OSKM) genes) can induce somatic cells to convert back to embryonic pluripotent stem cells^4,5^. Despite the simplicity of the process, it is typically inefficient and a-synchronized with less than 0.1-15% of the somatic donor cells undergo reprogramming over a period of 2-4 weeks^6^. In 2013, our lab has found that controlled and partial reduction of a key component of Mbd3/NuRD (Nucleosome Remodeling and Deacetylation) complex, named Mbd3, in concert with optimized OKSM delivery and naïve pluripotency conditions, can lead to highly efficient and rapid iPSC formation (up to 100% reprogramming efficiency within 8 days, in genetically controlled constellations) (Rais et al, 2013^1^). For murine iPSC induction, naïve pluripotency conditions consisted of controlled delivery of 2i/LIF, where 2i is applied 48 hours after reprogramming starts, 5% O2 hypoxia conditions, and Vitamin-C-containing Knock-out serum replacement (FBS-free and KSR only conditions - applied 48 hours after reprogramming initiation)^1^. We concluded that Mbd3/NuRD is a repressor of induction of pluripotency, from mouse EpiSCs, PGCs and somatic cells, as well as from human secondary *in vitro* differentiated fibroblasts.

Notably, another independent study by Grummt and colleagues^7^ showed that over expression of Mbd3 in MEFs blocks reprogramming, and its depletion promotes reprogramming of MEFs and partially reprogrammed cells. Further, previous work^8^ (including Dr. Jose Silva, Cambridge University, UK), has shown that the pluripotency factor Zfp281 directly recruits Mbd3/NuRD to repress Nanog promoter activity, and that inhibition of Zfp281 led to more than 3 fold increase in iPSC formation efficiency, thus supporting a repressive role for Mbd3/NuRD in iPSC formation. Notably, the fact that our genetically controlled Mbd3 depletion led to a radically more pronounced effect than Zfp281 depletion^1,8^, supports our suggested mechanism that Mbd3/NuRD may be acting more cardinally and upstream of Zfp281 in reprogramming regulation, by directly interacting with many other critical pluripotency promoting factors including OSKM (and likely other pluripotency factors)^1^.

Recently, a paper by Dos Santo et al.^2^ claimed that Mbd3 depletion has no influence on MEF reprogramming and yet has a negative effect on EpiSC and pre-iPSC conversion. Another non-peer reviewed communication by the same group (Bertone et al, 2015^3^) raised arguments aiming at challenging the validity of our iPSC efficiency quantifications, gene expression analysis, and the role of Mbd3/NuRD as an inhibitor for iPSC formation as presented in Rais et al. Nature 2013^1^. Here we address, reinterpret and challenge these studies, and provide arguments supporting our previous conclusions that indeed the somatic cells in our systems undergo rapid and authentic reprogramming as a result of Mbd3/NuRD depletion.

## Results

### Mbd3 transcript and protein levels in Mbd3^flox/-^ cell lines

Bertone et al.^3^ used our previously published gene array data sets^1^ and claimed that Mbd3 transcript level was only ∼20% depleted in Mbd3 flox/- cells and only ∼34% reduced in Mbd3 null cells. The gene array data sets published in our original study^1^ and used by Bertone et al.^3^ for making this claim, were harvested from cells that were expanded on irradiated WT mouse embryonic feeder cells (MEFs), and thus they are inappropriate for drawing conclusions regarding accurate Mbd3 transcription levels (Figure 1A). RT-PCR on cells expanded in feeder free conditions confirmed approximately 50% reduction of Mbd3 transcript in Mbd3^flox/-^ ESCs as expected (Figure 1B). Further, we now provide RNA-seq analysis that also confirms 50% reduction in Mbd3 transcript level in Mbd3^flox/-^vs. Mbd3^+/+^ WT cells (Figure 1C-D).

**Figure 1.**
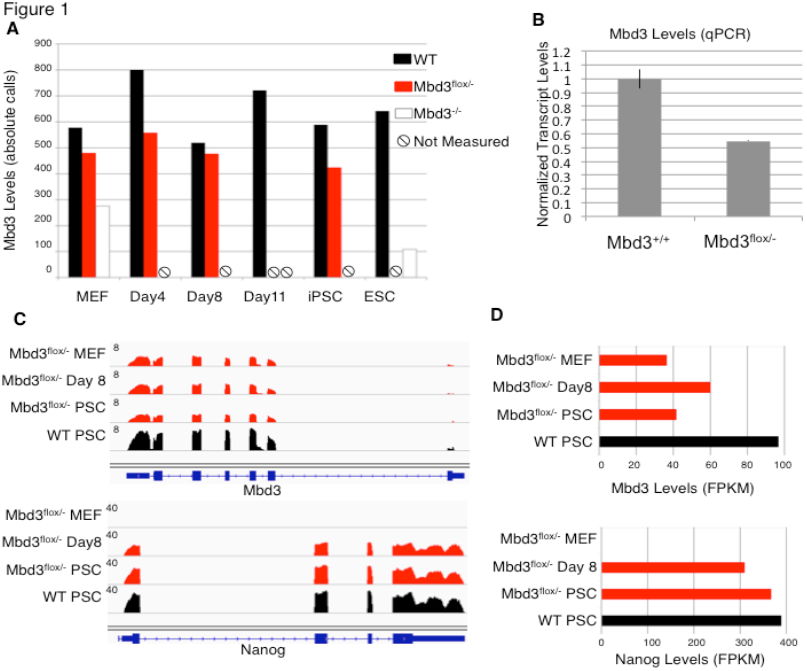
Measuring Mbd3 transcription levels. **A)** Mbd3 transcription levels measured by gene microarray datasets in Rais et al.^1^ Absolute Affymetrix calls are presented (after RMA normalization). Conditions in which expression levels were not measured are marked. Only the samples in A were harvested from cells grown on WT irradiated MEF cells. **B)** Mbd3 transcription levels measured by qPCR. **C)** Mbd3 and Nanog transcription landscape measured by RNA-Seq before, during and after reprogramming in Mbd3^fiox/-^ cell line as well as in Mbd3^+/+^ pluripotent stem cells. **D)** Normalized transcription levels (FPKM) of Mbd3 and Nanog, estimated from RNA-Seq data.

We showed in our paper (<u>Figure 1c, Rais et al Nature 2013</u>^1^) that Mbd3 protein level in the Mbd3^flox/-^ mouse ESCs were reduced up to 80% in comparison WT ESC. We have validated these results on 3 independent targeted Mbd3^flox/-^ ES clones (kindly provided by Brian Hendrich lab^9^) and expanded in <u>Serum(FBS)/LIF</u> conditions - 3flox18 clone (used in Rais et al. Nature 2013^1^), 3flox4 ES clone and 2lox30 clone (Figure 2A). Dos Santos et al.^2^ claimed absence of change in protein levels however they expand their ES cells in 2i/LIF conditions (Dos Santos et al. Cell Stem Cell 2014, Figure S7), which we note as a stabilizer for Mbd3 protein expression that reduces the differences between Mbd3^flox/-^ and WT cells (Figure 2B). <u>Please note that in Rais et al.</u>^1^ <u>we had to provide 2i after 48 hours, indicating that low Mbd3 levels in the fist 48 hours appears to be critical for this dramatic effect we described in iPSC generation from Mbd3^flox/-^ cells.</u> Unfortunately, Western blots for Mbd3 in FBS/LIF grown ESCs were not shown by the Dos Santos et al, but only 2i/LIF (Dos Santos et al. Cell Stem Cell 2014, Figure S7^2^). Further, our Western blot analysis confirms lower Mbd3 protein expression in Mbd3 ^flox/-^ MEFs before (Figure 2C) and after (Figure 2D) OSKM induction in comparison to WT cells.

**Figure 2.**
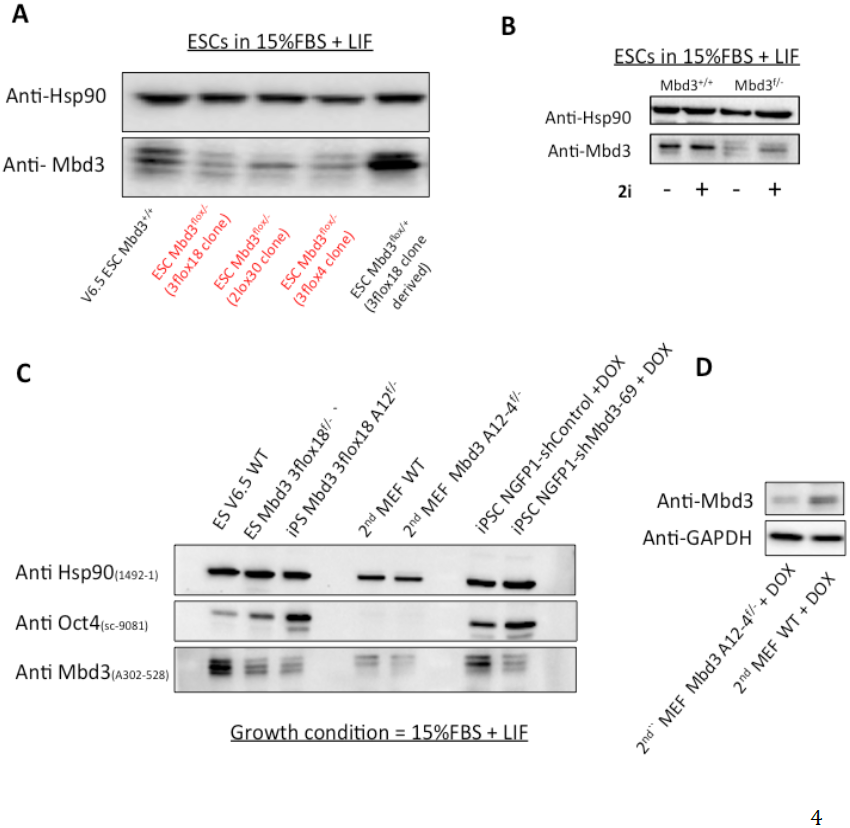
Mbd3 protein levels. **A)** Western blot analysis for Mbd3 in mouse ES cells expanded in FBS/LIF feeder free conditions. Three independently generated Mbd3^flox/-^ ES lines were used (3flox18, 3flox4, 2lox30), and 2 WT ES clones. **B)** Western blot analysis for WT or Mbd3^flox/-^ mouse ES cells expanded in FBS/LIF feeder free conditions with or without 2i (as indicated). **C)** Western blot analysis for Mbd3 and Oct4 in WT or Mbd3^flox/-^ cell samples. D) Western blot analysis on secondary WT and Mbd3^flox/-^ MEFs after 48 hours of OSKM (DOX) induction.

### Genomic analysis of iPSC reprogramming in Mbd3 WT and flox/- cells

In Rais et al.^1^, gene expression levels during reprogramming were measured in Mbd3^flox/-^ cells at days 0,4,8 after doxycycline (DOX) induction of OSKM, and compared to established iPS and ES cell lines. In addition, histone mark profiles and DNA methylation from the same samples were measured and compared. Importantly, these samples were harvested without selecting for any pluripotency markers or reporters. We showed that on day 8, the transcriptome and epigenome state in Mbd3^flox/-^ was indistinguishable from the ESC and iPSC reference samples, as shown by correlation and unsupervised clustering methods (<u>Rais et al, Fig 3a</u> and <u>Extended Data Fig 5a)</u>^1^. We now provided additional figures (Figure 3 and 4) that show clustering of the same previously published histone mark profiles^1^, corroborating the data presented in Rais et al.^1^ <u>Extended Data Fig 5a</u>. Here too, only Mbd3^flox/-^ on day 8, but not WT samples, is clustered with ESC/iPSC samples.

**Figure 3.**
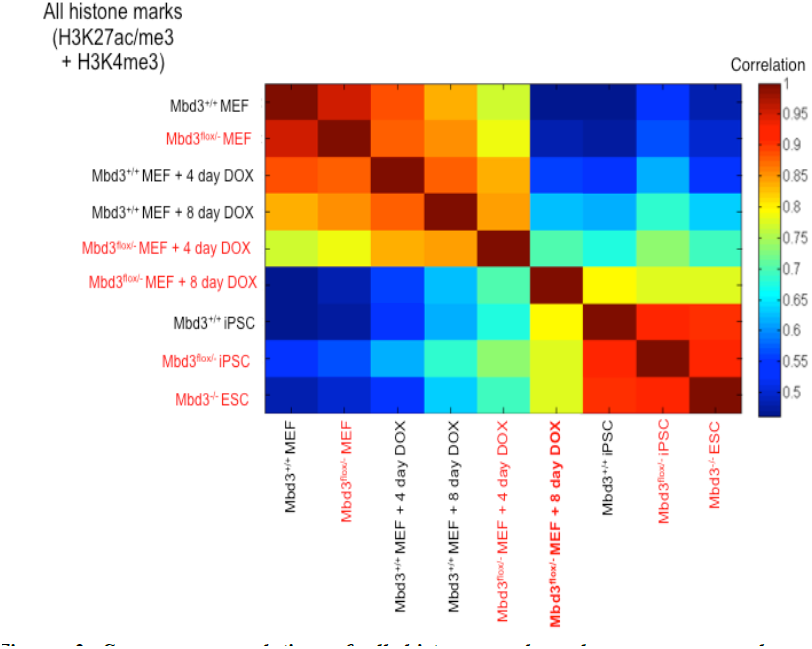
Spearman correlation of all histone marks values, as measured over differentially expressed genes. Chromatin IP-Seq was performed in donor fibroblasts before and after DOX induction and compared to established pluripotent iPSCs and ESC lines^1^. Gene profiles of H3K4me3, H3K27me3 and H3K27ac marks were extracted and normalized to z-scores. The value of each gene and each histone mark was chosen as the max value of the appropriate z-score profile (see Methods). Spearman correlation between the vectors described above (1541 genes with all histone marks.) By day 8, Mbd3^flox/-^ cells show epigenetic signature that strongly correlates with established ESCs and iPSCs line. Mbd3^+/+^ population does not show strong correlation with pluripotent ESCs-iPSC lines even after 8 days of reprogramming.

**Figure 4.**
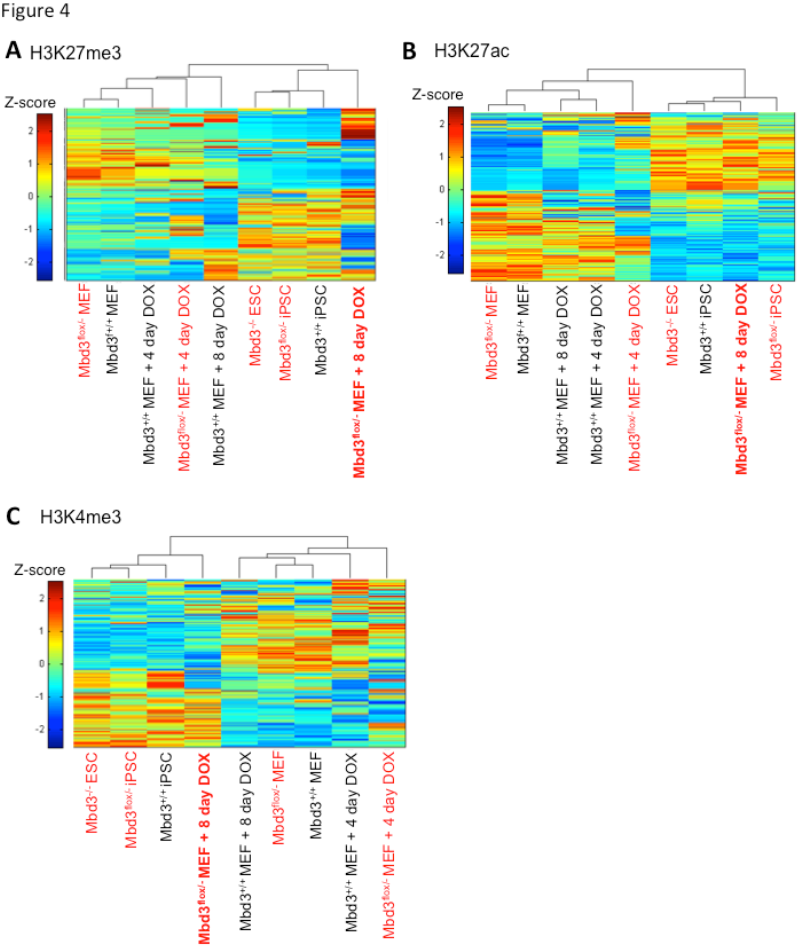
Clustering of each histone mark separately. Hierarchical clustering of standardized histone mark levels measured in 1541 differentially expressed genes between wild type MEF and ES samples (see Methods). Clustering was calculated using Spearman correlation as a distance metric and average linkage. H3K27me3, H3K27ac and H3K4me3 are presented (A,B,C, respectively). Results show that even when considering each histone mark separately, by day 8 Mbd3^flox/-^, but not MTbd3^+/+^, were epigenetically similar to established ESC and iPSC lines.

**Figure 5.**
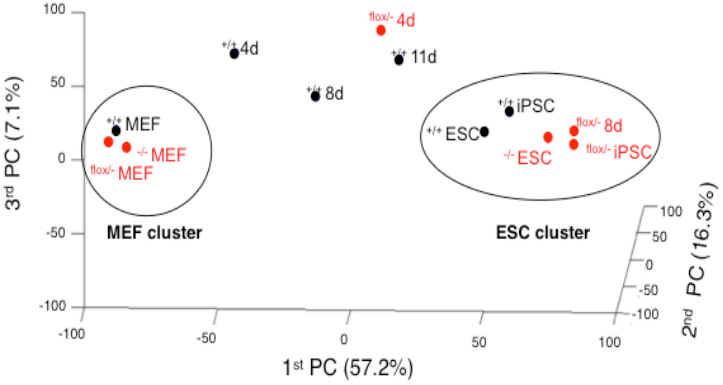
Transcriptional reprogramming dynamics in Mbd3^flox/-^ cells. Principle Component Analysis of transcriptional profiles measured in Mbd3^flox/-^ (red) and Mbd^flox/-^ (black) cell lines before, during and after reprogramming. PCA analysis finds two clusters for MEF cells andfor ES iPS cells (circles).

Transcription levels during reprogramming were also measured in Mbd3^+/+^ at days 0,4,8 and 11. WT Day 11 sample was selected for most comparisons as these cells had the advantage over day 8 Mbd3^flox/-^ cells, because they were reprogrammed for a longer period (**<u>which further allow WT reprogrammed cells take over at the expense of non-reprogrammed MEFs since the former proliferate almost twice faster</u>**). Still however, unsupervised hierarchical clustering (<u>Rais et al, Figure 3a</u>)^1^ showed that Mbd3^+/+^ samples, including day 8 and 11, do not cluster with ES and iPS control samples.

Bertone et al^3^ raised a concern that Mbd3^+/+^ day-11 seems to be indistinguishable from the Mbd3^flox/-^ day-8 sample, and that the difference between the samples may be dominated by biological noise. To test this claim we decided to recheck our analysis. We first tested the clustering of the samples using an independent method (Principal Component Analysis) over all active genes (n=17800), resulting in similar results (Figure 5), where only Mbd3^flox/-^ day-8 is clustered with ESC/iPSC samples, while WT day-11 and day-8 samples are in-between the clusters, and both do not cluster with ESCs or fully reprogrammed iPSCs.

When looking specifically at pluripotency promoting genes (Figure 6), we saw an elevated expression (at least 20% higher) of 60% of the genes in the Mbd3^flox/-^ compared to WT day-11. Fourteen genes were up regulated by more than 2 fold compared to WT day-11, including Rex1/Zfp42, Prdm14, Esrrb and Dppa4. This elevation was overlooked by Bertone et al^3^, due to their choice to show standardized log expression values, which reduces the signal differences. In Figure 6, the expression levels of pluripotent genes are presented in standardized log_2_ expression (left) and as fold-change compared to WT iPSC (right). In addition, we observed a number of somatic genes down regulated by more than 4 fold in Mbd3^flox/-^ compared to WT day-11, including Runx1, Thy1, Fgfr2 and Tgfbi, indicating of a more rapid shut-down of the somatic cellular program in Mbd3^flox/-^ compared to WT.

**Figure 6.**
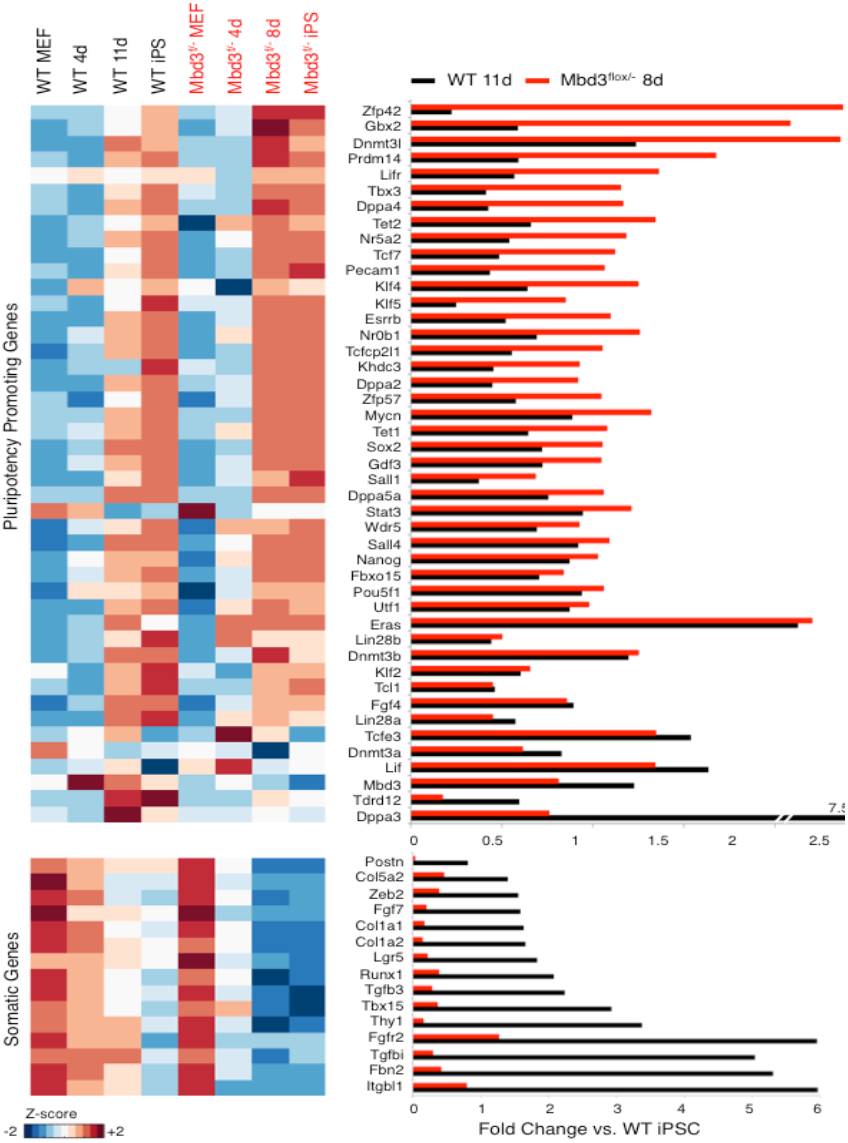
Expression pattern of pluripotency and somatic genes. Left: Expression pattern (Z-score transformed log_2_ expression) of pluripotency promoting genes (upper panel) and selected somatic genes (bottom panel). Right: Expression fold change compared to WT iPSC.

To make this analysis more general we selected genes that are differentially expressed between MEF and ESC/iPSC (t-test FDR <5% and > 4 fold-change), and used them to cluster all the samples (Figure 7A). This analysis yielded 583 up-regulated and 958 down-regulated genes (Table S1). Hierarchical clustering over these genes showed again that Mbd3^flox/-^ day-8 is highly similar to ESC/iPSC samples, while WT Mbd3^+/+^ samples cluster outside the ESC/iPSC cluster. Up regulated genes are enriched (p<10^-10^, FDR<5%) for cell cycle and stem cell maintenance genes (Figure 7B), and downregulated genes are enriched for developmental categories. This shows that the described differences appear to have biological meaning that cannot be regarded as biological noise. We next tested the dynamic change in the expression of these differential genes during reprogramming, and found that the changes were more rapid in Mbd3^flox/-^ cells compared to Mbd3^+/+^ cells (Figure 7C), both in up-regulated and in down-regulated genes.

**Figure 7.**
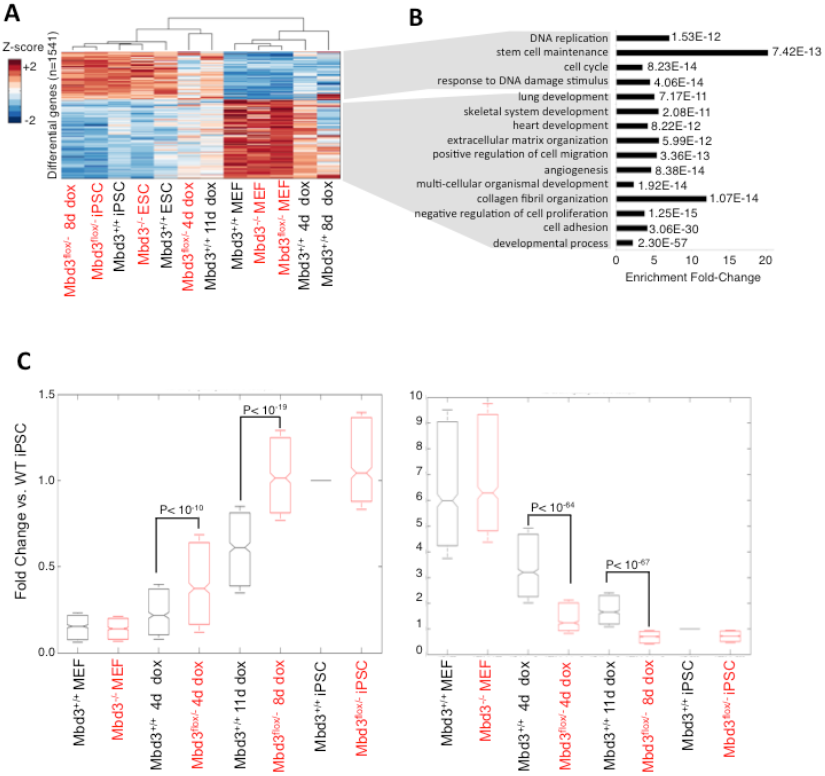
Differential gene signatures. **A)** Hierarchical clustering of all samples, over differentially expressed genes (between MEF and ESCZiPSC). **B)** GO function categories enriched in the two gene signatures. Bars indicate enrichment fold change compared to background (consisting of 19,682 Entrez genes), and numbers indicate Fisher exact test p-values (FDR<0.1%). **C)** Distribution of fold change (compared to WT iPSC) of genes signatures. Left: for genes that are up-regulated in ESC/iPSC (n=583) Right: for genes which are down-regulated in ESC/iPSC (n=958). Numbers indicate t-test p-values.

The results described above are consistent with similar analyses done with multiple histone marks (Figures 3-4, <u>Rais et al, Extended Data Figure 5A</u>)^1^ and DNA-methylation (<u>Rais et al, Extended Data Figure 5B-C</u>)^1^, reconfirming that Mbd3^flox/-^ day-8 is indistinguishable from the ES and iPS reference samples, but not WT samples. Unfortunately, these compelling evidences were ignored by Bertone et al^3^. Taken together, these different approaches show that both at the transcriptional and chromatin levels, the reprogramming of Mbd3^flox/-^ cells is authentic and more rapid than in WT (even when allowed to reprogram for additional days until harvesting at day 11).

### Use of un-matched reporters in WT and Mbd3^flox /-^ cells used for genomic analysis

Our original study^1^ entailed generation of over 20 independent clonal series carrying either GOF18 ΔPE-Oct4-GFP transgenic reporter (Addgene plasmid #52382) or complete GOF18 Oct4-GFP transgenic reporter (Addgene plasmid #60527), both valid reporters for murine naïve pluripotency and validated for specificity (e.g. <u>in Figure 2B of Rais et al</u>. we show 1 representative WT clone, and 2 independent representative flox/- clones). The clonal series selected for genomic analysis included an Mbd3^+/+^ clone that carries GOF18 ΔPE-Oct4-GFP transgenic reporter, and Mbd3^flox/-^ and Mbd3^-/-^ cells that carry GOF18 Oct4-GFP transgenic reporter (complete Oct4 enhancer region with DE and PE elements). These reporters can be identified when analyzing the Oct4 locus in genomic DNA input datasets and shown in Figure 1 by Bertone et al^3^. While in retrospect, it would have been more optimal to use cells carrying the matched reporters for the genomic analysis (as done for iPSC efficiency experiments throughout the Rais et al.^1^ manuscript), this does not affect the genomic results in any way. **Please note that we do not use Oct4-GFP or any other selection for sorting cells prior to conducting genomic experiments**. Thus, the difference in transgene reporters cannot influence the interpretation of our genomic analysis data in any way. Further, in these genomics studies, the endogenous Nanog and Oct4 loci are not manipulated and are identical between all cell lines as the Oct4-GFP reporters were introduced via random transgenesis and validated for specificity. Notably, in ground state naive 2i/LIF medium used for this analysis, the PE element is anyways not functionally relevant and thus the reporters cannot differentially influence the molecular outcome of any of our experiments. This is now clearly indicated in the GEO submission and in a Corrigendum (*Nature – submitted*) accompanying Rais et al.^1^.

**In summary, for the genomic analysis we harvested polyclonal donor cell cultures, without any selection or sorting, therefore the Oct4 reporters are completely irrelevant for those experiments.**

### Reporters used for iPSC efficiency quantification

For quantitative reprogramming efficiency assays eventually presented in Rais et al. Nature 2013^1^ used complete GOF18 Oct4-GFP transgene reporter and/or Nanog-GFP knock in reporters, both valid and accurate reporters for Naíve pluripotency acquisitions (as acknowledged by Bertone and co.^3^).

Further, the following comparative sets were used throughout <u>Rais et al.</u>^1^ study to establish the role of Mbd3/NuRD dependent changes:

1. EpiSC reversion in Figure 1b was conducted by using matched comparison for Nanog-GFP knock in for Mbd3 WT and flox/- cells.
2. In Figure 1f, we presented PGC reversion efficiencies form WT and flox/- based on Oct4-GFP complete GOF18 transgene coupled with direct staining for anti-Nanog immunofluorescence.
3. In Figure 2b and e^1^ –; we used randomly selected clones.(2 shown from flox/- and 1 for WT cells, out of over 20 clones generated and showed similar results) that carry full Oct4-GFP reporter both in WT and flox/- cells. Mbd3 rescue experiment in Figure 5e^1^ was conducted in the same Mbd3^flox/-^ line with the same complete Oct4-GFP transgene as delineated in Extended Data Figure 3a, and shows Mbd3 dependent phenotype.
4. In Figure 2d^1^ we used Nanog-GFP knock in together with staining for anti-Oct4 (after DOX removal) on WT and Mbd3 depleted samples (matched comparison on both types of cells as indicated in the Y-axis label).
5. In Figure 4^1^ we used shRNA Mbd3 secondary depletion on NGFP1 iPSC line carrying Nanog-GFP knock in reporter (matched comparison in control and KD cell line).
6. Notably, reprogramming progression can be estimated from the video analysis without the use of GFP reporter, for example by colony formation (which is dependent only on phase contrast or mCherry marker). Here too, colony formation was found to be more rapid in Mbd3^flox/-^ compared to WT cells (Figure 8, <u>Rais et al. Supplementary Movies</u>)^1^.

**Figure 8.**
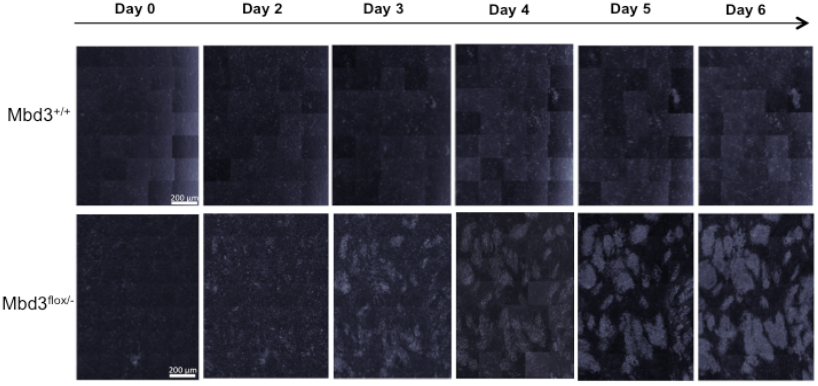
Reprogramming dynamics in Mbd3^flox/-^ cells. Phase contrast live imaging of colony formation during OKSM mediated reprogramming of Mbd3^flox/-^ (bottom) and Mbd^+/+^ (up) cell lines as previously shown in Rais et al.^1^.

In Rais et al. ^1^, live imaging was conducted on lines carrying complete Oct4-GFP full reporter transgene (matched between both cell lines), but not Nanog Knock-in. because these assay are conducted on entire single wells and 5X magnification, and only Oct4-GFP complete reporter is strong enough to allow detection with this low magnification (in our hands, Nanog-GFP is weak for detection with this low magnification, but is reliable for FACS analysis or manual quantitation in 20X magnification). Notably, ΔPE-Oct4-GFP WT and flox/- reporter lines yield identical results to full Oct4-GFP transgenic reporter, but we regarded this as trivial since we used 2i/LIF conditions. In fact, ΔPE-Oct4-GFP transgene reactivation kinetics reported in WT and C/EBPa transgenic Pre-B cell reprogramming in <u>Figure 2a</u> and <u>Extended Data Figure</u> 4E by Di Stefano et al.^10^ to be very similar to our WT and Mbd3^flox-^ cells, respectively, harboring complete GOF18 Oct4-GFP transgenic reporter^10^.

In summary, reprogramming quantitation was used by multiple quantification methods in different systems to yield a similar and valid conclusion.

### Specificity and effect of integration sites of Oct4-GFP transgenic reporter

Generally speaking, the use of genetic knock-in reporters and transgenic models with defined insertions is more favorable. As such, we have also used Nanog-GFP knock-in reporter system throughout our manuscript^1^ (but not for live imaging for the reason indicated above).

However, the complimentary use of secondary systems carrying integrations is very common and valid in iPSC mechanistic studies (e.g. O’Malley et al. Nature 2013 ^11^ by Keisuke Kaji and colleagues– Piggyback transposition system for examples insert very high integration levels – often over 20). Further Dos Santos et al. Cell Stem Cell 2014^2^ used transgenic Oct4-GFP reporter without mapping insertions:

*“EpiSCs: Mbd3fl/– and Mbd3–/– EpiSCs were derived from ESCs as previously described (Guo et al., 2009). Briefly, ESCs transfected with pPB-EOS-GFPires-Puro (EOS-GiP; GFP-ires-Puro under the control of early transposon promoter and Oct4 and Sox2 enhancers) were cultured in Fgf2/Act. A medium for at least 10 passages before analysis. To obtain a pure EpiSC culture, GFP+ cells were removed by FACS ” (A direct quote form their methods section)*.

While we do not map integration sites in the secondary lines used^1^, we overcome such potential biases by 1) using multiple secondary lines (e.g. Rais et al. Figure 2b)^1^, **2) by rescue reconstitution of Mbd3 WT and mutant alleles (Rais et al. Figure 5e)**^1^, 3) by using a distinct NGFP-KD system that does not carry any Oct4-GFP transgene integrations (Rais et al. Figure 4)^1^, 4) by conducting EpiSC reprogramming on lines which carried only defined knock-in allele and no random integrations (Rais et al. Figure 1B), 5) and by boosting (although not up to 100% efficiency in this setting) reprogramming via non-integrating Mbd3 or Chd4 siRNA (Rais et al. Fig 1a. and Rais et al. Extended Data Fig 8b)^1^. These massive complimentary approaches prove dependence on Mbd3/NuRD perturbation to yield the observed phenotypes in our study.

Finally, the claim by Bertone and colleagues^3^ that the use of complete Oct4-GFP reporter is less optimal than ΔPE-Oct4-GFP (regardless whether introduced as a transgene or knock-in), since the former is expressed in other lineages like the hypoblast, is misleading. ΔPE-Oct4-GFP is not exclusively expressed in naïve pluripotent cells either, but also in other Oct4+ lineages like PGCs^12^. Thus, based on the latter, one can reach the conclusion that none of the Oct4 reporters is appropriate iPS efficiency measurement. We refute these claims, and emphasize that, particularly when 2i/Lif conditions are used, both reporters are valid and specific for naïve mouse iPSC formation^12^. We reemphasize anyways that we used matched reporter for quantitative iPSC analysis and defined Nanog-GFP knock in reporters, but provide the previous discussion for the sake of scientific argumentation (“even if” scenario).

### Complete depletion of Mbd3 blocks somatic cell proliferation and viability

In Rais et al. Nature 2013^1^ clearly indicated that near 100% complete inhibition of Mbd3 in somatic cells or EPiSCs results leads to abrupt decrease in cell proliferation and viability. As such we greatly focused on using hypomorphic Mbd3^flox/-^ cell lines and compared them to Mbd3^+/+^ WT cells. In cases where we achieved high reprogramming efficiency of Mbd3^-/-^ cells (Rais et al, Figure1-2, Extended Data Figure 3B,C)^1^, we always started with Mbd3^flox/-^ cells (and NOT Mbd3^-/-^ cells as used in Dos Santos et al.) in order <u>to ensure low and residual Mbd3 protein levels already during the first 48 hours of OSKM induction.</u> Afterwards, 2i was supplemented with tamoxifen to achieve complete ablation of Mbd3. This is a critical point that was highlighted in our paper:

*“Mbd3^-/-^ MEFs (but not pluripotent cells) experience accelerated senescence and proliferation capacity loss, and thus Mbd3^-/-^ somatic cells were reprogrammed by applying tamoxifen on <u>Mbd3^flox/-^</u> cells only after 48 h of OSKM induction ”. (A direct quote from Rais et al. Nature 2013^1^ Methods section)*

The results presented by Dos Santos et al.^2^ systematically used different conditions where they did either one of the following:

Dos Santos et al.^2^ used Mbd3^-/-^ cells that were maintained for multiple passages as null cells and only afterwards OSKM were introduced. As we had previously indicated1, these cells dramatically lose their proliferation capacity, and indeed all growth proliferations curves shown in Dos Santos et al. validate this result (even though they were carried for 4 days only). **Therefore the entire findings by Dos Santos et al.^2^ claiming that Mbd3 facilitates epigenetic reprogramming can be trivially explained by inducing cell proliferation block, and in our opinion, has nothing to do per se with epigenetic reprogramming. Further, the latter decrease in cell proliferation occurs even without OSKM induction, further supporting the notion that the inhibition of reprogramming results from simply hampering cell proliferation in donor cells, rather than epigenetic reprogramming per se.** The Mbd3 “rescue” by Dos Santos et al. ^2^, rescued also cell proliferation and hence reprogramming. Further, the levels of Mbd3 have not reached Mbd3 levels in WT cells, and were actually the same or even lower than those shown in their Mbd3^flox/-^ samples. Throughout our work, we avoided generating Mbd3^-/-^ in such a manner as described by Dos Santos et al. ^2^ because these cells do not robustly proliferate and thus do not reprogram, which, in our opinion, is very trivial.
For Tamoxifen induced deletion experiments, Dos Santos et al.^2^ used Mbd3^flox/flox^ cells (and not Mbd3^flox/-^ cells), and applied tamoxifen at different time points. However, these experiments lead to complete deletion of Mbd3, and not during the critical window of early reprogramming, which we highlighted as critical^1^.
Finally, Dos Santos et al.^2^ did not compare WT vs. Mbd3^flox/-^ cells as starting somatic donors for reprogramming, which is the most relevant comparison and the most robust comparative one presented in our paper^1^.

In summary, our findings in fact are consistent with Dos Santos et al^2^, as they use conditions that ensure either complete inhibition of Mbd3 and block cell proliferation, or avoid optimal depletion of NuRD activity during a critical early reprogramming window, which we previously highlighted^1^ as critical determinants for capturing the beneficial effect of Mbd3/NuRD depletion on iPSC reprogramming.

## Summarizing Conclusions

In summary, we disagree with Bertone, Silva and colleagues^2,3^, and stress that all efficiency sets presented in our Rais et al.^1^ study are valid and comparable, and live up to widely applied standards in the reprogramming field. Mbd3/NuRD is a major pathway that inhibits the maintenance and induction of pluripotency^1,7^ and its controlled manipulation should be an integral pathway for inducing and maintaining naïve pluripotency in a variety of species. Along with the importance of the observations regarding the inhibitory role of Mbd3 in the reprogramming process^1,7^, it is clear that the we still has a long way to go in order to fully understand the mechanisms underlying Mbd3 functions (including NuRD dependent and independent ones).

**As such, in our opinion, the relevant next challenge will be to identify signaling pathway inhibitors that lead to controlled Mbd3 depletion and/or find genetic alternative ways to inhibit Mbd3/NuRD repressive activity (i) early in the reprogramming process <u>and</u> (ii) without blocking somatic cell proliferation and viability.** Framing the latter challenges, will allow routine and widespread adoption of inhibiting this potent repressive pathway in iPSC reprogramming towards naïve pluripotency, and possibly from multiple species.

## Acknowledgements

J.H.H is supported by a generous gift from Ilana and Pascal Mantoux; the New York Stem Cell Foundation, FAMRI, the Kimmel Innovator Research Award, the ERC (StG- 2011-281906), the Leona M. and Harry B. Helmsley Charitable Trust, BIRAX initiative, The Sir Charles Clore Research Prize, the Israel Science Foundation (Morasha, NFSC, ICORE and Regular research programs), the ICRF Foundation, Helen and Martin Kimmel Institute for Stem Cell research (HMKISCR) the Benoziyo Endowment fund, Fritz Thyssen Stiftung, Erica A. and Robert Drake. J.H.H. is a New York Stem Cell Foundation - Robertson Investigator. We thank Weizmann Institute management for providing critical financial and infrastructural support. We thank Prof. Naama Barkai for discussions and comments.

## Supplementary Table Legend

**Table S1: Differentially expressed genes between MEF and ESC/iPSC samples.**

## Methods

### Poly-A RNA sequencing

RNA was extracted from Trizol pellets, and utilized for RNA-seq by TruSeq RNA Sample Preparation Kit v2 (*Illumina*) according to manufacturer’s instruction. DNA sequencing was conducted on Illumina Hiseq1500.

Total RNA was extracted from the indicated cell cultures using PerfectPure RNA cultured cell kit (cat#2302340, 5 Prime). To avoid DNA contaminations all samples were treated with DNase (5 Prime). RNA integrity was evaluated on Bioanalyzer (Agilent 2100 Bioanalyzer), requiring a minimal RNA integrity number (RIN) of 8.5. Libraries were prepared according to Illumina's instructions accompanying the TruSeq RNA Sample Preparation Kit v2 (RS-122-2001). Sequencing was carried out on Illumina HiSeq2500 according to the manufacturer’s instructions, using 10pM template per sample for cluster generation, and sequencing kit V2 (Illumina).

### RNA-Seq Analysis

Poly-A RNA sequencing was measured in mRNA extracted from Mbd3^flox/-^ MEFs, days 8 after Dox (OKSM) induction, established Mbd3^flox/-^ iPS cells and ES cells, as well as in Mbd3^+/+^ V6.5 ES cells. The paired-end reads were aligned to mouse genome version mm10 with TopHat2 aligner (v2.0.8b), using TopHat2 default input parameters. Transcriptional profiles are visualized using IGV v2.3. FPKM levels (Fragments-per-kilobase-per-million reads) were estimated using Cufflinks package with “-p 3 –u” parameters, and GTF file downloaded from ensemble (version GRCm38.74). This newly generated RNA-Seq dataset will be made publically available upon publication of a new follow-up study from our group by Zviran et al. (manuscript in preparation).

### DNA microarray Analysis

Microarray data were published previously^1^: nine samples (Mbd3^flox/-^ MEF, Mbd3^flox/-^ day-4, Mbd3^flox/-^ iPSC, Mbd3^-/-^ MEF, Mbd3^-/-^ ESC, WT MEF, WT day-4, WT day-11 and WT iPSC) were published in Rais et al, 2013^1^ and two samples (WT ESC and WT day-8) were previously published by our group (GSE35775) (Mansour et al. Nature 2012).

CEL files of all samples were analyzed with Matlab affyRMA command, with MoGene_1_0-st-v1.r3.cdf annotation file, resulting in expression levels of 35513 Affymetrix probe sets. Probe sets without any call above 5 (in log_2_ scale) were filtered out, and the left probes were translated to genes. For genes with several probe sets, the one with the highest average expression (across all samples) was chosen as the representative expression pattern. This procedure resulted in 17800 active genes that were further analyzed. PCA analysis was carried out with 17800 active genes, using princomp Matlab command (version R2011b). To generate expression heatmap of pluripotent and somatic genes (Figure 6), log_2_ expression values were transformed to z-score ((x- *μ*)/ *σ*), and presented with Matlab (R2011a) HeatMap command.

Differentially expressed genes between MEF samples (WT MEF, Mbd3^flox/-^ MEF, Mbd3^-/-^ MEF) and ESC/iPSC samples (WT ESC, WT iPSC, Mbd3^flox/-^ iPSC and Mbd3^-/-^

ESC) were chosen using t-test, FDR <5% and above > 4 fold-change Hierarchical clustering of these genes was generated by Matlab clustergram command, using Spearman correlation as a distance metric and average linkage, and per-gene standardization to z-score. Box plot describing distribution of fold change (compared to WT iPSC) was generated using Matlab boxplot command, without outliers.

### Functional Enrichment

Enrichment of GO process categories was calculated using Fisher exact test. GO categories which passed p-value < 10^-10^ (FDR<0.1%) are presented along their enrichment fold change and p-values.

### Histone mark profiles

Chip-Seq data were previously conducted and published^1^ (NCBI GEO accession number GSE49766). Histone mark profiles were calculated using in-house script. Shortly, this script generates a matrix of read densities in given genomic intervals. In this case, the profiles of all 29,952 Entrez genes (mm9, taken from UCSC known gene tables) were calculated between 1kb upstream to TSS and TES. These read densities were then converted to z-score by normalizing each position by the mean and standard deviation of the sample noise 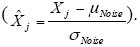 Noise parameters were estimated for each sample from 6*10^7^ random bp across the genome. In the histone mark correlation and clustering analysis each gene and each histone mark is represented with the maximal z-score measured in the profile of that gene, where the profiles were calculated as described above. Clustering of histone marks was carried out on concatenated vectors that include all marks for every gene in tandem. Hierarchical clustering of these chromatin marks was generated by Matlab clustergram command using Spearman correlation as a distance metric and average linkage.

### Mouse stem cell lines and cell culture.

WT or mutant Mbd3 mouse ESC clones were expanded in mouse ES medium consisting of: in 500ml DMEM (Invitrogen), 15% Fetal Bovine Serum (Biological Industries), 1mM glutamine (Invitrogen), 1% nonessential amino acids (Invitrogen), 0.1mM *β*-mercaptoethanol (Sigma), penicillin-streptomycin (Invitrogen), 10 *μ* g recombinant human LIF (Peprotech). Stem cell lines deficient for Mbd3 were obtained as previously described (Mbd3^+/+^, Mbd3^flox/-^, Mbd3^flox/+^ and Mbd3^-/-^). Mycoplasma detection test are weekly conducted, to ensure exclusion of any contaminated cells. MEFs were obtained from E12.5 embryos as previously described^1^, and expanded in mouse ESC medium with or without Dox as indicated.

### Western blot analysis

Western blot analysis on samples harvested from feeder free conditions, was performed by using the following primary antibodies: anti-Mbd3 (1:1000, A302-528, Bethyl), anti- OCT4 (1:1000, sc-9081, Santa Cruz), anti-Gapdh (1:5000 Epitomics 2251-1), anti-Hsp90 (1:1000 Epitomics 1492-1).

### qPCR analysis

Total RNA was isolated using the RNeasy Kit (Qiagen). 3 *μ* g of total RNA was treated with DNase I to remove potential contamination of genomic DNA using a DNA Free RNA kit (Zymo Research). 1 *μ* g of DNase-I-treated RNA was reverse transcribed using a First Strand Synthesis kit (Invitrogen) and ultimately re-suspended in 100 *μ*l of water. Quantitative PCR analysis was performed in triplicate using 1/50 of the reverse transcription reaction on Viia7 platform (Applied Biosystems). Error bars indicate standard deviation of triplicate measurements for each measurement. Primer Sequences: Gapdh: For –5’-CATTGTGGAAGGGCTCATGACCA-3’, Rev- 5’- GCAGGGATGATGTTCTGGGCAG-3’; Mbd3: For-5’- GGCCACAGGGATGTCTTTTACTATAG -3’, Rev- 5’-GTTGTGGCTTGCTGCGG-3’.

